# Initial leads to combat streptogramin resistance generated from X-ray fragment screening against VatD

**DOI:** 10.1101/2025.01.31.635826

**Authors:** Pooja Asthana, Sonya Lee, Christian M. MacDonald, Ian Seiple, James S Fraser

## Abstract

Streptogramins are potent antibiotics targeting bacterial ribosome. The synergistic binding of group A and B streptogramins to 50S-ribosome, yields bactericidal effects. However, their efficacy is compromised by resistance mechanisms, including enzymatic acetylation of group A streptogramins by Virginiamycin Acetyl Transferase (Vat) enzymes, which reduces their affinity for ribosomes. Using fragment-based drug discovery we identified starting points for development of VatD inhibitors. X-ray crystallography screening revealed three primary fragment-binding sites on VatD. In the acetyl binding subsite, fragments stabilized distinct conformational states in critical residues, His82 and Trp121. In the antibiotic binding site, two fragments formed interactions that could be leveraged for competitive inhibition. Elaborations of these fragments showed weak inhibition of VatD activity, indicating potential for further optimization. These findings establish initial hits that could restore streptogramin efficacy by targeting VatD directly, providing a structural foundation for inhibitor development against resistant bacterial strains.

## Introduction

The relentless evolution of antibiotic resistance poses an imminent threat to public health, necessitating the development of new chemical compounds to combat bacterial infections ^1^. Streptogramins are a potent class of antibiotics that are used to treat a range of bacterial infections including the vancomycin-resistant *Staphylococcus aureus* (VRSA), vancomycin-resistant enterococci (VRE) ^2^. In addition to their clinical applications, streptogramins have also been used as growth promoters in animal agriculture ^3^.

Streptogramin antibiotics consist of structurally unrelated group A (e.g. dalfopristin) and group B (e.g. quinupristin) compounds. Like many clinically-relevant antibiotics, streptogramins target the bacterial ribosome, with group A streptogramins binding to the peptidyl transferase center (PTC) and group B streptogramins binding adjacently at the nascent peptide exit tunnel (NPET) ^4,5^. Group A streptogramins, along with other PTC-binding antibiotics like chloramphenicol and lincosamides, interfere with tRNA substrate positioning and catalytic peptidyl-transfer ^6,7^. NPET-targeting antibiotics, such as macrolides and group B streptogramins, extend into the NPET and inhibit elongation and ribosome translocation ^5,7,8^ (**Figure 1A**). Streptogramins A and B have evolved to bind synergistically in the 50S ribosomal subunit with a complementary molecular interface that enables a synergistic, bactericidal effect in some organisms ^9^. Individually, however, these compounds exhibit only bacteriostatic activity ^4,5,9^.

**Figure 1:**
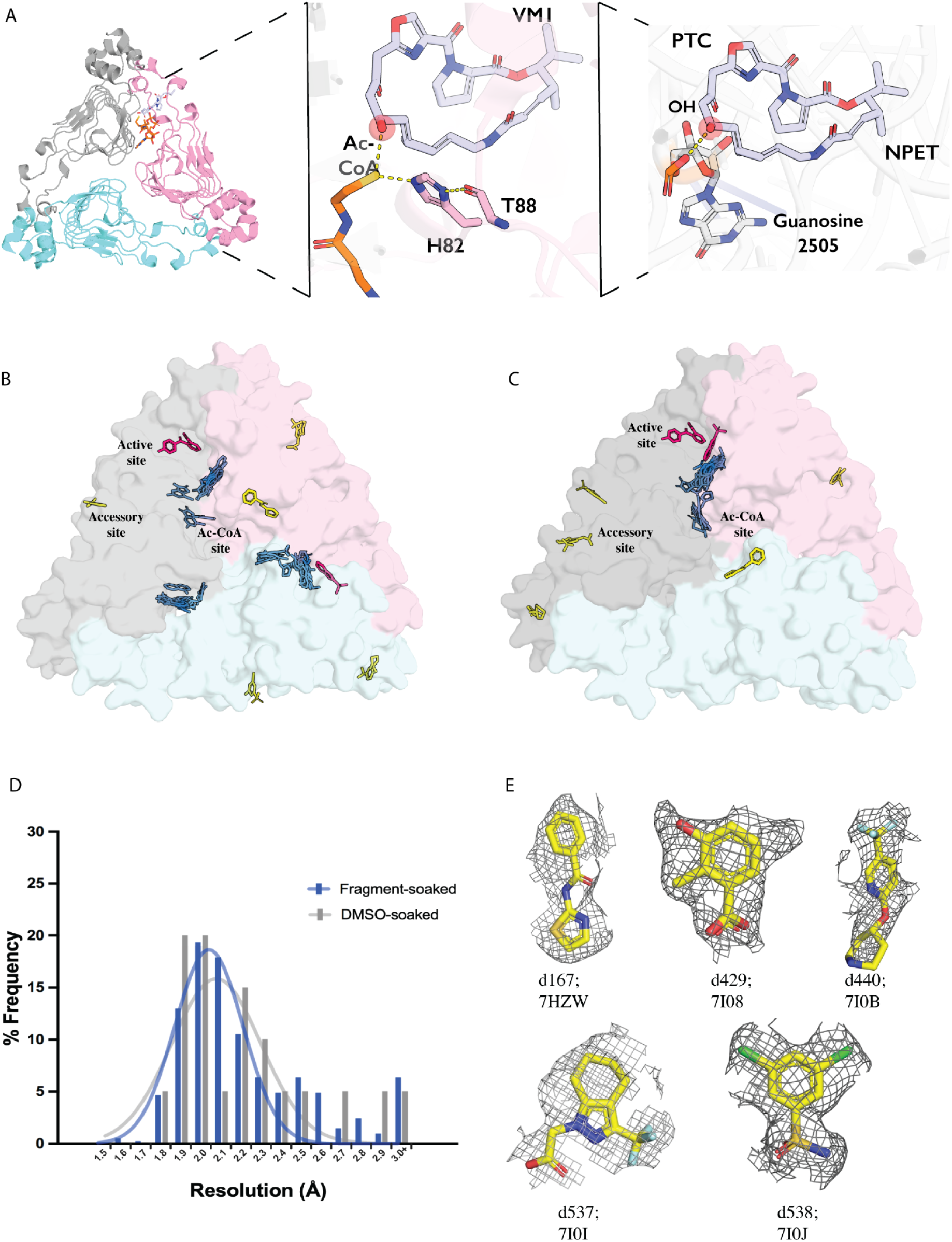
A) Schematic representation of VatD inhibiting binding of streptogramin antibiotic class A, virginiamycin M1 to the bacterial (*E. coli*) 50S ribosomal subunit (PDB: 4U25). The ribosome is shown as a cartoon representation in grey. Ac-CoA (PDB: 1KK4) and VM-1 are shown in stick representation in orange and purple, respectively. The OH group of VM-1 which gets acetylated is highlighted in red sphere. VatD is shown in cartoon representation with three chains in grey, pink, and blue. B) Unaligned crystal structures of VatD complexed with fragments. C) Crystal structures of VatD complexed with fragments aligned to VatD-apo as the reference. The fragments are colored by their presence in the Antibiotic binding site (pink), Ac-CoA site (blue), and non-active sites (yellow). D) Histogram showing the resolution range of VatD crystal soaked with 10% DMSO (grey) and 10% fragments (blue) for 4 hours. The X-axis represents the resolution range, Y-axis represents the percentage of frequency. E) Gallery of density for the non-active/surface site fragments: d167 (7HZW), d429 (7I08), d440 ()7I0B, d537 (7I0I), d538 (7I0J). The event map density shown is at sigma 1.5 contour.

Bacteria have developed various adaptive mechanisms to overcome protein synthesis inhibition by streptogramins. Similar to other PTC-targeting antibiotics, the resistance to group A streptogramins can be mediated by target site modifications. Most notably, the activity of the Cfr rRNA methylases modifies A2503 of the 23S RNA directly blocking the binding of many antibiotics, including group A streptogramins, to PTC ^10,11^. Additionally resistance can arise from the binding of ABC-F proteins that utilize ATP hydrolysis to dislodge antibiotics, including streptogramins, from the PTC ^12–14^. The ABC-F proteins insert into the E-site of ribosome, triggering diverse conformational changes that facilitate drug release ^14^.

While Cfr and ABC-F mechanisms impact diverse antibiotics that bind in the PTC, there is also a key resistance mechanism specific to group A streptogramins mediated by a class of enzymes known as <u>V</u>irginiamycin <u>A</u>cetyl <u>T</u>ransferases (Vats). Vats confer resistance by transferring an acetyl group from the co-substrate acetyl-CoA (Ac-CoA) to the free hydroxyl group of the O18 of group A streptogramins (Sugantino and Roderick 2002; Kehoe et al. 2003; Stogios et al. 2014) (**Figure 1A**). When bound in the ribosome, the O18 hydroxyl group forms a hydrogen bond with the phosphate oxygen of G2505 and acetylation results in the disruption of this hydrogen bond, which significantly decreases the affinity of group A streptogramins for PTC ^8^.

Vat enzymes are encoded by plasmid borne genes *vat*(A), *vat*(B), and *vat*(C) from *Staphylococcus aureus*, *vat*(D)/*sat*(A), *vat*(E)/*sat*(G), and *vat*(H) from *Enterococcus faecium*. The Vat enzymes belong to a superfamily of trimeric enzymes characterized by a left-handed β-helical (LβH) fold built from a repeating hexad amino acid sequence motif ^15^. This LβH fold is closely related to the fold of xenobiotic acetyltransferases (XAT), such as chloramphenicol acetyltransferase (CAT) ^16^. The trimeric Vat has three catalytic centres present at each of the interchain interfaces ^15^. Each centre has one site for the binding of Ac-CoA and one antibiotic binding site (for the streptogramin A co-substrate). A conserved histidine residue in Vats (H82) has been identified as a critical catalytic base required for acetylation ^15^. While previously we made modifications at different positions of the group A scaffold to overcome the resistance caused by Vats ^17^, directly inhibiting the Vat protein itself is an attractive strategy for potentiating group A streptogramin antibiotics. However, there are currently no compounds that inhibit the Vats directly. The strategy of developing inhibitors to enzymes that deactivate antibiotics has been spectacularly successful for β-lactams, with β-lactamase inhibitors now commonly used in the clinic ^18^.

In this study, we focus on VatD from *E. faecium* as a model to identify potential inhibitors of Vat-mediated resistance. We applied X-ray crystallography based fragment screening to efficiently explore a relatively broad chemical space to discover novel small-molecule fragments binding to VatD. X-ray based screening is a good fit for identification of VatD inhibitors because it is unclear which of the antibiotic or Ac-CoA binding sites on Vat is best suited for the development of a competitive inhibitor. Our findings reveal distinct fragment-binding sites on the surface of VatD and alternative receptor conformations stabilized by hits binding to the acetyl site. Furthermore, we created analogs of the fragments binding at the antibiotic interaction site, which act as weak competitive inhibitors of streptogramin A inactivation. Collectively, this study establishes a foundation for the development of direct Vat enzyme inhibitors, opening up new avenues for combating streptogramin resistance.

## Results

### Fragments bind to co-factor, substrate, and surface sites of VatD

To enable crystallographic fragment screening against VatD, we first optimized reproducible crystallization and DMSO-soaking conditions modeled after the procedures created at the X-CHEM facility ^19^. Initially, we identified a new crystal form grown in 0.2 M NH4Cl, and 17% Peg 3350 that diffracted to 2.0-2.8 Å resolution (**Supplementary Figure 1**). However, when these crystals were treated with 5% DMSO, they showed ice rings and sporadic twinning issues. We were unable to overcome these challenges and therefore identified an alternative system (0.1 M Tris, 25% Peg 1000, pH 8.5) that was robust to DMSO up to 10% for 5 hours (**Figure 1D**). Using these conditions, we soaked VatD crystals with 411 fragments, comprising the Enamine Essential (320 fragments) and a small custom library (91 fragments), for 4-6 hours before harvesting and vitrification. We collected X-ray diffraction data from these crystals with most crystals diffracting to 2.0-2.5 Å resolution (**Figure 1D**).

To identify fragments that bound to VatD, we analyzed the resulting electron density maps using PanDDA ^20^. This analysis identifies “events” as localized deviations in fragment-soaked electron density maps from the background DMSO-soaked maps. Across our 411 soaks, we discovered 226 events dispersed across 65 sites. We visually inspected all events due to the relatively modest resolution in some of our datasets. After examining the maps, we identified 27 “true positive” fragments, with 5 fragments bound in multiple locations in the same crystal (**Figure 1B**, Table 1). The asymmetric unit of the crystal is a trimer, which is the biochemically relevant oligomeric state of VatD. This crystal form, therefore, presents each potential binding site in 3 slightly different crystallographic environments. While most fragments bound only in a single chain in the asymmetric unit, 2 fragments bound in the same location across all three chains of VatD (**Figure 1B**, Table 1).

**Table 1:** All of the fragments hits obtained from soaking of Enamine_320 and UCSF_91 library.

|  | Dataset | No of copies | Site | PDB ID |
| --- | --- | --- | --- | --- |
| 1 | d516 | 1 | Antibiotic |  |
| 2 | d521 | 1 | Antibiotic |  |
| 3 | d193 | 3 | Ac |  |
| 4 | d232 | 1 | Ac |  |
| 5 | d287 | 1 | Ac |  |
| 6 | d374 | 1 | Ac |  |
| 7 | d384 | 1 | Ac |  |
| 8 | d398 | 1 | Ac |  |
| 9 | d474 | 1 | Ac |  |
| 10 | d499 | 1 | Ac |  |
| 11 | d501 | 2 | Ac |  |
| 12 | d519 | 1 | Ac |  |
| 13 | d397 | 1 | CoA |  |
| 14 | d229 | 2 | CoA |  |
| 15 | d275 | 1 | CoA |  |
| 16 | d330 | 1 | CoA |  |
| 17 | d425 | 2 | CoA |  |
| 18 | d436 | 3 | CoA |  |
| 19 | d438 | 1 | CoA |  |
| 20 | d545 | 1 | CoA |  |
| 21 | d560 | 1 | CoA |  |
| 22 | d116 | 1 | CoA |  |
| 23 | d429 | 1 | accessory |  |
| 24 | d440 | 1 | accessory |  |
| 25 | d537 | 1 | accessory |  |
| 26 | d538 | 1 | accessory |  |
| 27 | d167 | 1 | accessory |  |

To analyze the receptor environment of the bound fragments, we aligned all three chains into a common reference apo-VatD (**Figure 1C**). From the aligned structures, we sorted the bound fragments into three major categories: 1) 20 fragments bound in the Acetyl-CoA binding site; 2) 2 bound in the antibiotic binding site; and 3) 5 bound at non-active sites scattered around the surface of VatD (**Figure 1C**). Despite having strong event map density, the 5 non-active site ligands did not have obvious paths to elaboration as allosteric modulators and were not analyzed further (**Figure 1E**).

### Fragments bound in the Ac-CoA site stabilize alternative conformations

VatD catalyzes the transfer of an acetyl group from Ac-CoA, which binds in a ∼24 Å long narrow channel that is formed at the interface of two chains of the VatD trimer (**Figure 2A**). Therefore, the Ac-CoA site is an attractive candidate for developing competitive inhibitors. The fragments that overlap with the Ac-CoA site are present in two main locations: largely overlapping with either the acetyl or CoA binding subsites, with 10 fragments bound in each location (**Figure 2B**).

**Figure 2:**
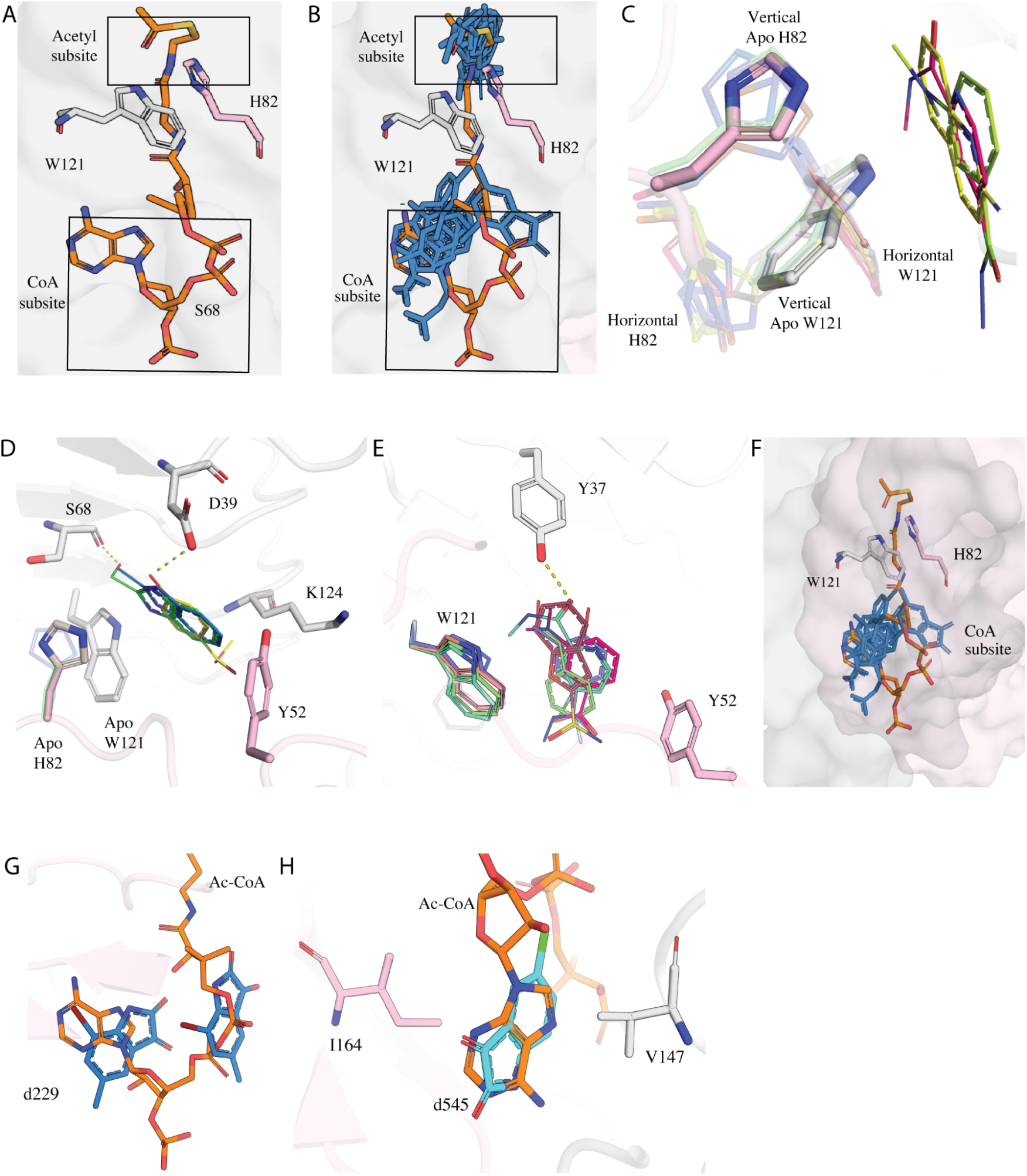
A) The subdivision of Acetyl-CoA site formed by the two chains of VatD trimer showing the two main subsites: Acetyl and CoA subsite. The two chains of apo VatD are shown in surface representation (grey, pink). H82 and Y121 residues emerging from distinct chains of the apo protein are shown in sticks. The Ac-CoA is shown in sticks in orange. B) The binding of fragments (blue) in the acetyl and CoA subsite. C) Fragments showing the horizontal and vertical conformations of H82 and W121 relative to fragments. The apo/vertical conformation is highlighted while the vertical confirmation is shown with less transparency. Fragments d193 (PDB: 7HZX), d232 (PDB: 7HZZ), and d287 (PDB: 7I01) are shown as sticks. D) E) Hydrophobic and hydrophilic interactions shown by fragments in the horizontal and vertical conformation. The H-bonds are represented by dashed lines. F) Fragments bound in the CoA subsite shown in stick representation in blue. G) Fragment d229 (PDB: 7HZY) in blue present in double occupancy overlapping at the adenosine moiety and pantothenic arm of Ac-CoA. H) Fragment d545 (PDB: 7I0K) in cyan showing a complete overlap with the adenosine. d545 (PDB: 7I0K) makes hydrophobic interactions with V146 and I164.

Fragments bound in the acetyl binding subsite were accommodated in two different receptor conformations characterized by distinct side chain rotamers populated by H82 and W121. The first conformation observed in four fragments (d232, d287, d501, and d519) resembles the apo state of VatD (**Figure 2C**). In this conformation, H82 and W121, maintain a stack in “vertical conformation” relative to the fragments. All four fragments make hydrophobic interactions with Y52, W121, and K124 (**Figure 2D**). In addition, fragments d232, and d501 make H-bond interactions with D39, while d519 shows a H-bond with S68 (**Figure 2D**).

In the second conformation, stabilized by 6 fragments (d193, d374, d384, d398, d474, and d499), H82 and W121 reorient to a “horizontal conformation” to accommodate the fragments (**Figure 2C**). In general, these fragments closely approach the apo H82 and W121 “vertical” conformations, likely leading to clashes and thus stabilizing the horizontal conformation. Similar to vertical conformers, these fragments also make hydrophobic interactions with Y52 and W121 (**Figure 2E**). d398 and d474 make additional H-bonds with Y37 (**Figure 2E**). The displacement of H82 is notable because it is the only absolutely conserved amino acid in the family and it acts as a catalytic base in the transfer of the acetyl group from CoA to the OH of antibiotic ^15,21^. The H82A mutation results in the reduction of acetylation activity by >10^5^ fold. This suggests that it would be unable to mutate for resistance and therefore interactions with this residue are important to consider in inhibitor design.

An additional ten fragments (d116, d229, d275, d330, d397, d425, d436, d438, d545, d560) bind within the CoA subsite (**Figure 2F**). These fragments form hydrophobic interactions with residues V147 and I164. Notably, fragment d229 is present in two copies in the CoA site, with one copy present overlapping with the location of the CoA adenosine moiety and the other copy overlapping with the location of the pantothenic arm (**Figure 2G**). Given the chemical composition of the fragment library used for soaking, it is unexpected that many fragments structurally similar to adenosine were not identified as hits, as we observed in another fragment campaign of an ADP-ribose binding macrodomain ^22^. Furthermore, even when fragments resembling adenosine, such as d397, were detected, they did not replicate the characteristic adenosine interactions. Only d545 has a relatively complete overlap with the adenosine and shows a similar pattern of hydrophobic interactions (**Figure 2H**). Collectively, the observations from the Ac-CoA binding site suggests that the 6 hits stabilizing the horizontal conformation are the most promising candidates for potential inhibitors.

### Fragments bound in the antibiotic binding site leverage distinct binding principles from streptogramins A

Out of all the fragment hits identified, only fragments d516 and d521 successfully found their way into the antibiotic site (**Figure 3** A,B). Both fragments were observed at a single active site within the trimeric VatD structure. The binding sites of d516 and d521 overlapped with the SA (VM-1) binding site, each fragment occupying distinct regions of the long alkyl chain of VM-1 (**Figure 3B**). Structural alignment of the fragment-bound forms to the apo structure revealed minor conformational changes in L108 and P109 side chains (**Supplementary Figure 7A**). Similar changes are also observed in a structure of VatD complexed with VM-1 and dalfopristin (PDB: 1KHR, 1MR9). Notably, the conformation of the catalytic residue H82 remained unchanged in both fragment-bound, SA bound, and apo forms.

**Figure 3:**
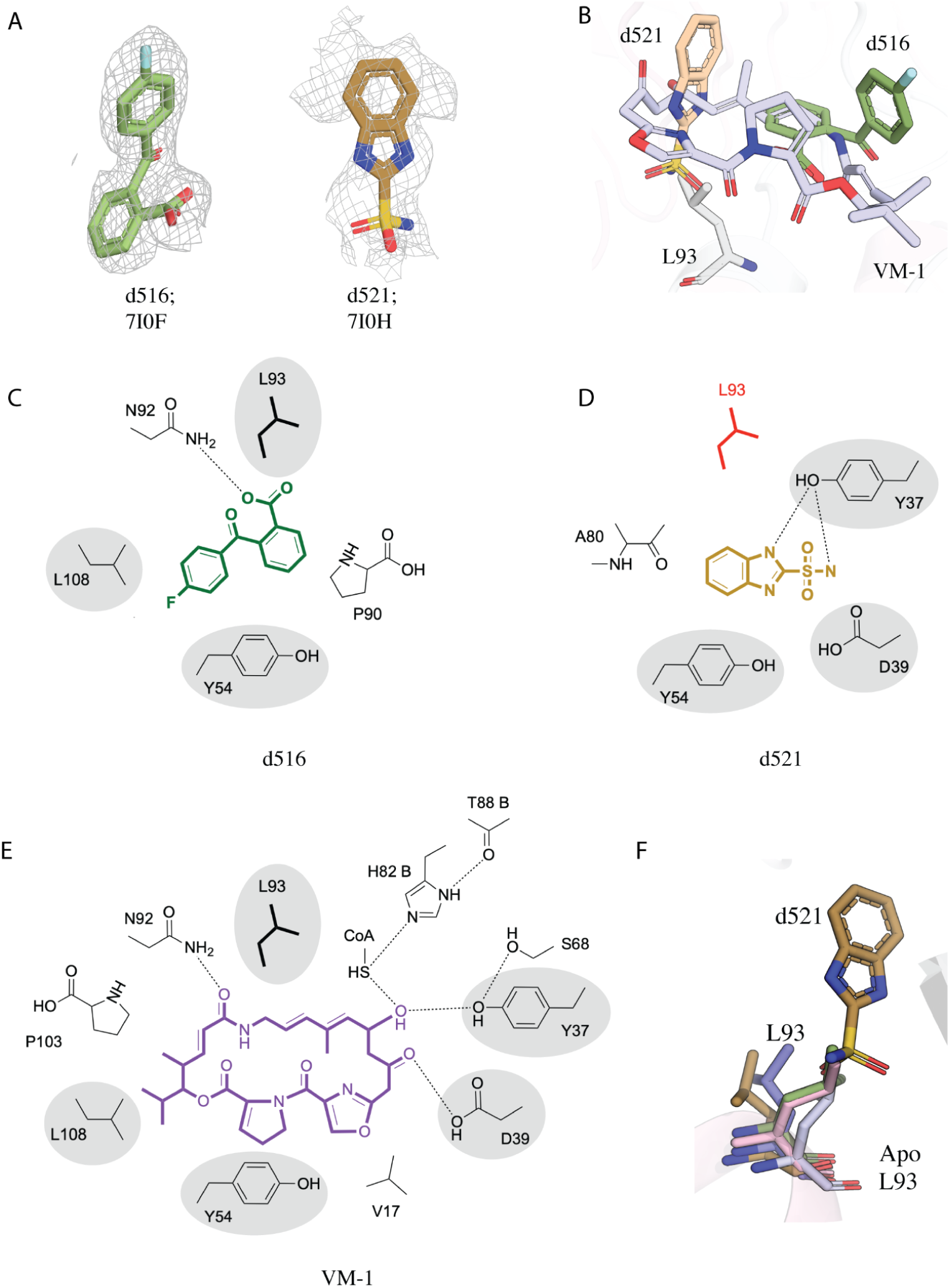
A) The event density map of fragments d516 (green, PDB: 7I0F) and d521 (brown, PDB: 7I0H) identified by PanDDA. B) Superposition of VatD complexed with d516 (green), d521 (brown), and VM-1 (purple, PDB: 1KHR) to the apo-VatD showing the binding site of fragments and SA. Apo-L93 is shown in sticks. C) D) E) The schematic diagram depicting the hydrophilic and hydrophobic interactions at the active site of VatD-d516 (green), VatD-d521 (brown), and VatD-VM-1 (purple) complexes. The hydrogen bonds are represented as dashed lines. Shared residues highlighted are in grey. F) The conformation flip of L93 side chain in d521 bound structure when compared to the apo-VatD (pink), VM-1-VatD (light purple, PDB: 1KHR), dalfopristin-VatD (dark purple, PBD: 1MRL).

Fragment d516 forms a single hydrogen bond (2.7 Å) between its carboxylate and the side chain of N92, which also interacts similarly with VM-1 (**Figure 3C**, E). In addition, d516 engages in multiple hydrophobic interactions with residues Y54, P90, L93, and L108 of VatD (**Figure 3C**). Similar hydrophobic interactions are also found in the SA bound structure as well except for P90 which is unique to d516 (**Figure 3E**).

In contrast, fragment d521 exhibited a hydrogen bond with Y37, similar to the interaction seen with VM-1. An additional hydrogen bond was observed between the amine group of d521 and D39 of VatD (**Figure 3D**). Furthermore, d521 formed hydrophobic interactions with A80 and Y52. The fragment would clash with the side chain of L93 in the apo (and 516-bound) conformation and consequently it flips to a state that more resembles the dalfopristin-bound structure (**Figure 3F**). Collectively, the observed interactions between these fragment hits and VatD recapitulate several contacts present in the SA-bound complex. This structural mimicry suggests that these fragments can serve as starting points for further elaboration.

### Elaborations at Acetyl-binding subsite did not yield potent compounds

Next, we wanted to optimize the compounds bound to the acetyl-binding subsite. We decided to focus on fragments that displaced the key catalytic residue H82 and then further narrowed our focus on two fragments with very strong electron density (d384 and d499) (**Figure 4A**). We overlaid these two compounds and explored merging their features. This generated analogs that grew along the vectors represented by the union of the two compounds. First, we built out from the two terminal methyl groups of d384. Second, we explored cyclization to add additional bulk in d384 in regions overlapping with the aromatic group of d499 and extending towards the antibiotic binding site. In considering which compounds to order, we prioritized trying to maintain the H-bond between the amino group of K124 and d384 while gaining interactions with Y37, D39, and S68 (**Figure 4B**, C).

**Figure 4:**
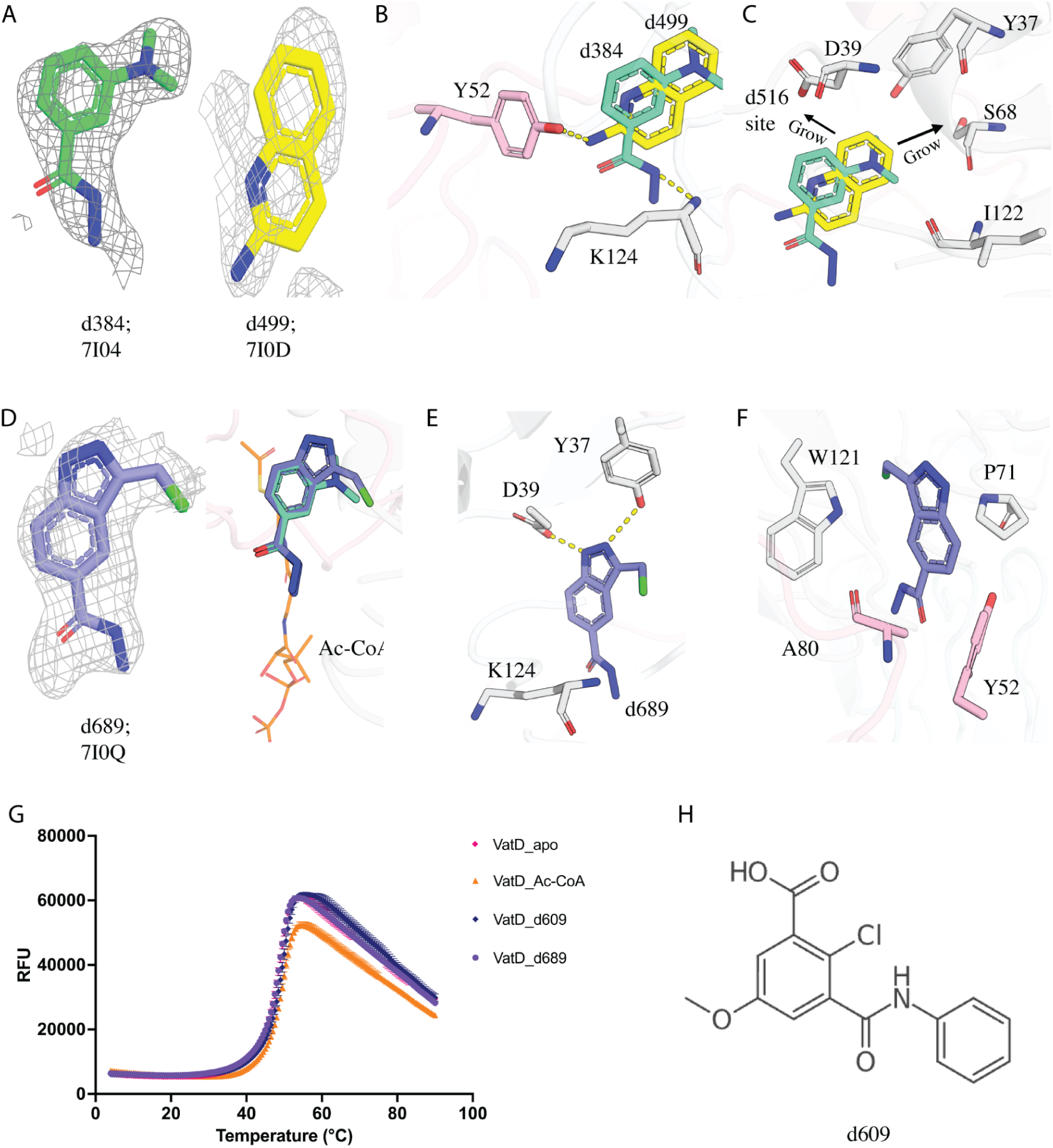
A) The event density map for fragment d384 (green, PDB: 7I04) and d499 (yellow, PDB: 7I0D) identified by PanDDA. B) The binding site of fragment d384 (green) and d499 (yellow) at the dimeric interface of VatD showing the H-bond between d384-K124, and d499-Y52. The residues coming from two different chains of VatD are shown in grey and pink color. C) The design principle of d384 and d499, extensions towards the d516/antibiotic binding site. D) The PanDDA event density map for fragment d689 (purple, PDB: 7I0Q). The binding of d689 in VatD showing the overlap with parent fragment and acetyl subsite (orange sticks). E) F) d689 showing new additional contacts formed with VatD residues. The H-bond is shown as dashed lines. G) The DSF curve showing the Tm for VatD bound to d609 (blue), d689 (purple), Ac-CoA (orange), Apo (pink). Ac-CoA is a positive control showing a change in Tm, however d689 does not show any Tm shift wrt to the Apo. H) The chemical structure of d609 which showed a Tm shift in DSF, but not found in PanDDA analysis.

We ordered 23 compounds (**Supplementary Figure 2**) to assay by differential scanning fluorimetry (DSF) and X-ray crystallography. From DSF, only one compound, d609, showed a temperature shift (Tm= 49.7 ℃) (Table 2, **Figure 4G**, H). However, we could not locate d609 in the electron density by PanDDA analysis from the X-ray data we collected from soaked crystals. We soaked the remaining 22 compounds into crystals of VatD and observed density only for one compound (d689, **Figure 4D**). Although this compound showed no shift by DSF, it is bound with strong overlap to the parent fragment d384 in the acetyl binding subsite (**Figure 4D**, G). In addition, d689 shows new H-bonds with Y37, and D39 (**Figure 4E**). Interestingly, these H-bonds are made by the atoms responsible for the additional cyclization beyond that present in d384. Additionally, hydrophobic contacts with Y52, P71, A80, and W121 (**Figure 4F**). Similar to the parent fragment (d384), d689 also shows a vertical conformation for H82 and W121 residues. However, consistent with the DSF results, this compound showed no effect on catalytic activity (**Supplementary Figure 5**). Despite the intriguing structural insights gained from d689, our attempts to elaborate the d384 hit into potent inhibitors were unsuccessful.

**Table 2:** List of compounds that showed binding in the DSF assay with their melting point (Tm) as calculated from the DSF world. The Tm for the control sample (without fragment) was 48.6 ℃.

| Compound | T<sub>m</sub><br>(°C) | SD | Binding<br>site |
| --- | --- | --- | --- |
| d706 | 48.57 | 0.21 | Antibiotic |
| d721 | 49.83 | 0.06 | Ac-CoA |
| d758 | 54.93 | 0.12 | Antibiotic |
| d705 | 50.90 | 0.1 | Ac-CoA |
| d724 | 52.45 | 0.07 | Ac-CoA |
| d689<br>(d384 ext) | 48.6 | 0 | Ac-CoA |
| Ac-CoA | 49.7 | 0.1 | Positive<br>control |
| VM-2 | 60.7 | 0 | Positive<br>control |
| d718 | 50.60 | 0.7 | No<br>structure |
| d763 | 49.50 | 0.0 | No<br>structure |
| d766 | 50.07 | 0.15 | No<br>structure |
| d711 | 49.57 | 0.06 | No<br>structure |
| d609<br>(d384 ext) | 49.70 | 0.1 | No<br>structure |

### Elaborations at the antibiotic binding site identify two promising series

In parallel, we designed analogs of the two fragments that bound, in non-overlapping positions, at the antibiotic binding site. The strategy behind the design and elaboration of d521 centered on two key approaches: (1) utilizing the antibiotic VM-1 as a guiding scaffold and (2) extending the d521 molecule linearly off the imidazole to probe potential interactions with residues within the active site (**Figure 5A**). These linear extensions were anticipated to establish interactions with residues N14 or L93, while maintaining the two hydrogen bond interactions we observed between the compound and VatD (**Figure 5A**). We ordered 17 compounds to assay by DSF and X-ray crystallography (**Supplementary Figure 3**). While none of these compounds stabilized VatD by DSF, soaking VatD crystals with these compounds identified one hit, d677 (**Figure 5B**). Surprisingly, this compound did not bind at the antibiotic site, but rather at the CoA subsite of Ac-CoA where it made hydrophobic contacts similar to other CoA subsite hits (**Figure 5C**, D). When compared to the parent fragment d521, it lacks the amino group at the sulfonamide moiety, which forms a hydrogen bond with Y37 and may therefore be crucial for active site binding. This compound did not inhibit the enzymatic activity of VatD.

**Figure 5:**
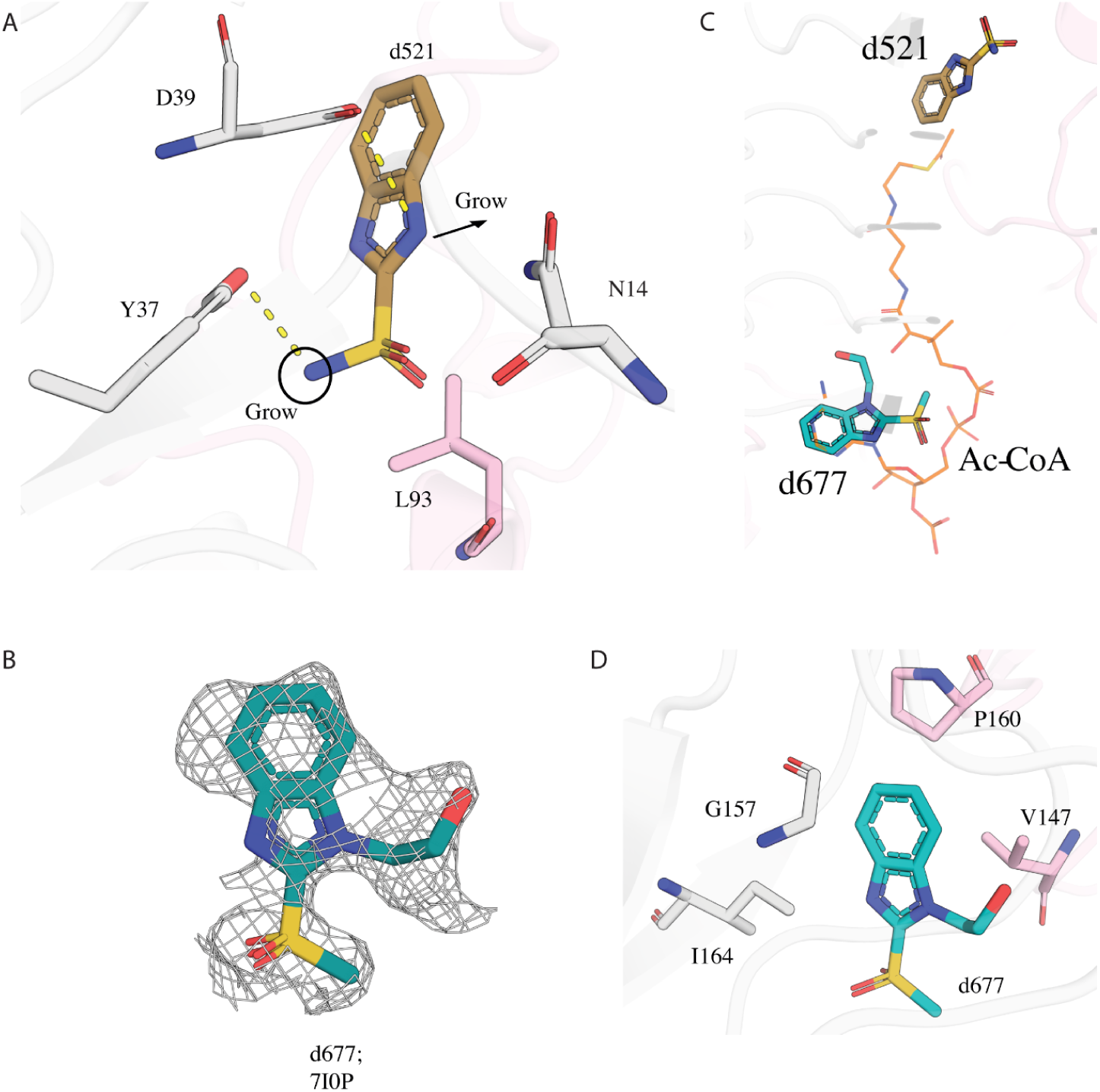
A) The design principle for elaboration of d521(brown, PDB:7I0H). The residues coming from two different chains of VatD are shown in grey and pink. B) PanDDA event density map of d677 in green (PDB: 7I0P). C) The presence of d677 at the acetyl subsite of Ac-CoA (PDB: 1KK4) in VatD. The Ac-CoA is shown in sticks in orange. The parent compound d521, binding in the antibiotic site is also shown in sticks in brown. D) The hydrophobic interactions made by d677 with the VatD residues.

For d516, our strategy focused on linear extensions from the carboxylic group, with the aim of picking up interactions similar to those observed with d521 with nearby Y37. These extensions were also informed by the structure of VM-1, leading to designs incorporating either pyrrolidine or oxazoline ring modifications. We also prioritized maintaining the H-bond between the carboxylic group of d516 and residue N92 (**Figure 6A**). We ordered 27 compounds to assay by DSF and X-ray crystallography (**Supplementary Figure 4**). From DSF, 4 compounds (d705, d721, d724, and d758) showed a temperature shift (**Figure 6C**). We soaked all 27 compounds into VatD crystals. We were surprised to find three compounds that did not generate a DSF shift (d619, d626, d666) bound in the CoA subsite (**Figure 6B**) and three of the compounds (d705, d721, d724) with significant DSF shifts bound in the acetyl subsite. Although the density for d721 was too weak to generate a model that unambiguously orients the molecule in the binding site, the density did support the overall location in the acetyl subsite. We assayed the three DSF and crystallography-positive compounds in the catalytic assay and found that only d721 and d724 showed inhibition activity. To test whether these compounds formed aggregates that can result in a non-specific enzyme inhibition ^23,24^, we incorporated a detergent (0.01% Triton X-100) into our catalytic assay. The results of the assay revealed Triton X-100 eliminated inhibition for d721 and d724 (**Figure 6G**), suggesting a non-specific aggregation-based inhibition mechanism.

**Figure 6:**
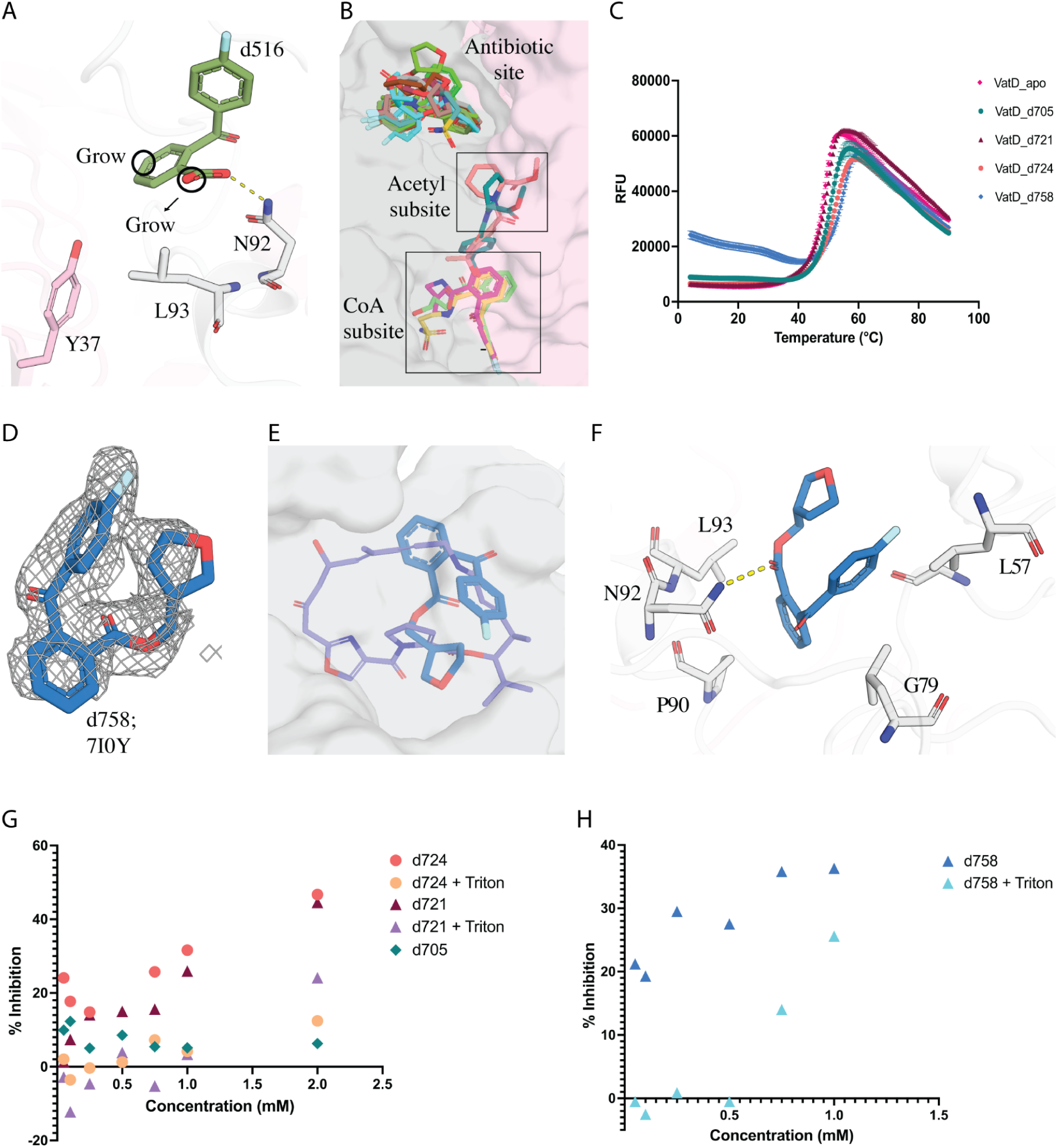
A) d516 (PDB: 7I0F) design principles for elaboration B) Distribution of 14 hits obtained from the analogs of d516 bound at 3 sites in VatD. C) The DSF curve showing the change in Tm for VatD bound to d705 (green, PDB: 7I0T), d721 (maroon red), d724 (orange, PDB: 7I0W), d758 (blue, PDB: 7I0Y), and apo (pink). D) d758 PanDDA event density map. E) Overlap of d758 (blue, PDB: 7I0Y) with VM-1 (purple, PDB: 1KHR) in sticks, and the VatD is shown in surface representation. F) d758 interactions with VatD residues G) The % inhibition graph of d705 (green), d721 (maroon red, purple), and d724 (orange, yellow) in absence and presence of 0.01% Triton-X in the assay. The concentration of the compounds range from 0.05 mM to 2 mM in the assay. H) The % inhibition graph of d758 in presence (blue) and absence (cyan blue) of 0.01% Triton-X in the assay. The concentration of the d754 ranges from 0.05 mM to 1 mM in the assay.

In addition, as anticipated, we observed 8 compounds bound in the antibiotic site (d698, d703, d706, d714, d756, d758, d764, d767, **Supplementary Figure 6A**). Most of these compounds exhibited significant structural overlap with the parent fragment d516, with the exception of d703, d706, and d756 (**Supplementary Figure 6B, C**). Notably, all compounds induced a conformational flip in residues L108 and P109, similar to that observed in the SA-bound structure (also seen in d516 and d521) when compared to the apo form (**Supplementary Figure 7B**). DSF screening revealed that while 7 compounds did not show thermal shifts, one compound d758 exhibited a significant shift of ∼6 ℃ (**Figure 6C**). This compound showed additional promising interactions with hydrophobic L57 and G79 that are also observed in VM-1 (**Figure 6E**, F). We assessed d758 for inhibition, and found strong inhibition at high concentrations. Addition of detergent (0.01% Triton X-100) to the assay showed a loss of inhibition suggesting it could be a non-specific inhibitor (**Figure 6H**). Despite the detergent sensitivity of d758, the initially promising DSF and X-ray results led us to continue exploring this series with another generation of designs.

### Further optimization of compounds bound to the antibiotic site identifies a weak inhibitor of VatD activity

We ordered 3 analogs of d758 designed to mimic chemical features present in VM-1 (**Figure 7A**). Although none of the compounds showed any change in melting temperature by DSF (**Figure 7C**), we discovered electron density for 2 of them, d797 and d798, in X-ray datasets of soaked crystals (**Figure 7B**). Surprisingly, the binding poses of these two compounds were quite different. While d798 overlaps with d758 (parent compound), with a slight shift in the placement of the furan ring (**Figure 7D**), d797 binds in a completely different pose within the antibiotic site. In d797, the furan ring was replaced with an oxazoline to more closely match VM-1. However, in the d797-bound structure this oxazoline ring is oriented in entirely the opposite direction of the design goal (**Supplementary Figure 8**). This compound binds in a hydrophobic pocket of VatD with key binding site residues being L57, P90, L93, P103, and P109 (**Figure 7E**). However, it did not maintain the H-bond with N92 as seen in the parent compound d758 and VM-1 (**Figure 7D**). This H-bond is lost due to the change in the binding pose of d797, placing the carbonyl oxygen away from N92. Despite the altered binding pose, d797 also shows a flip in the L108 and P109 side chains similar to other fragments we identified (**Supplementary Figure 7B**). We assayed these three compounds in the catalytic assay and found that only d797 inhibited catalytic activity. Unlike the previous compounds, this inhibition was not detergent-sensitive, giving more confidence in the specificity of the inhibition mechanism (**Figure 7F**). The estimated IC50 is greater than 500 µM, suggesting that this is a weak inhibitor, but a valuable starting point for more focused future efforts.

**Figure 7:**
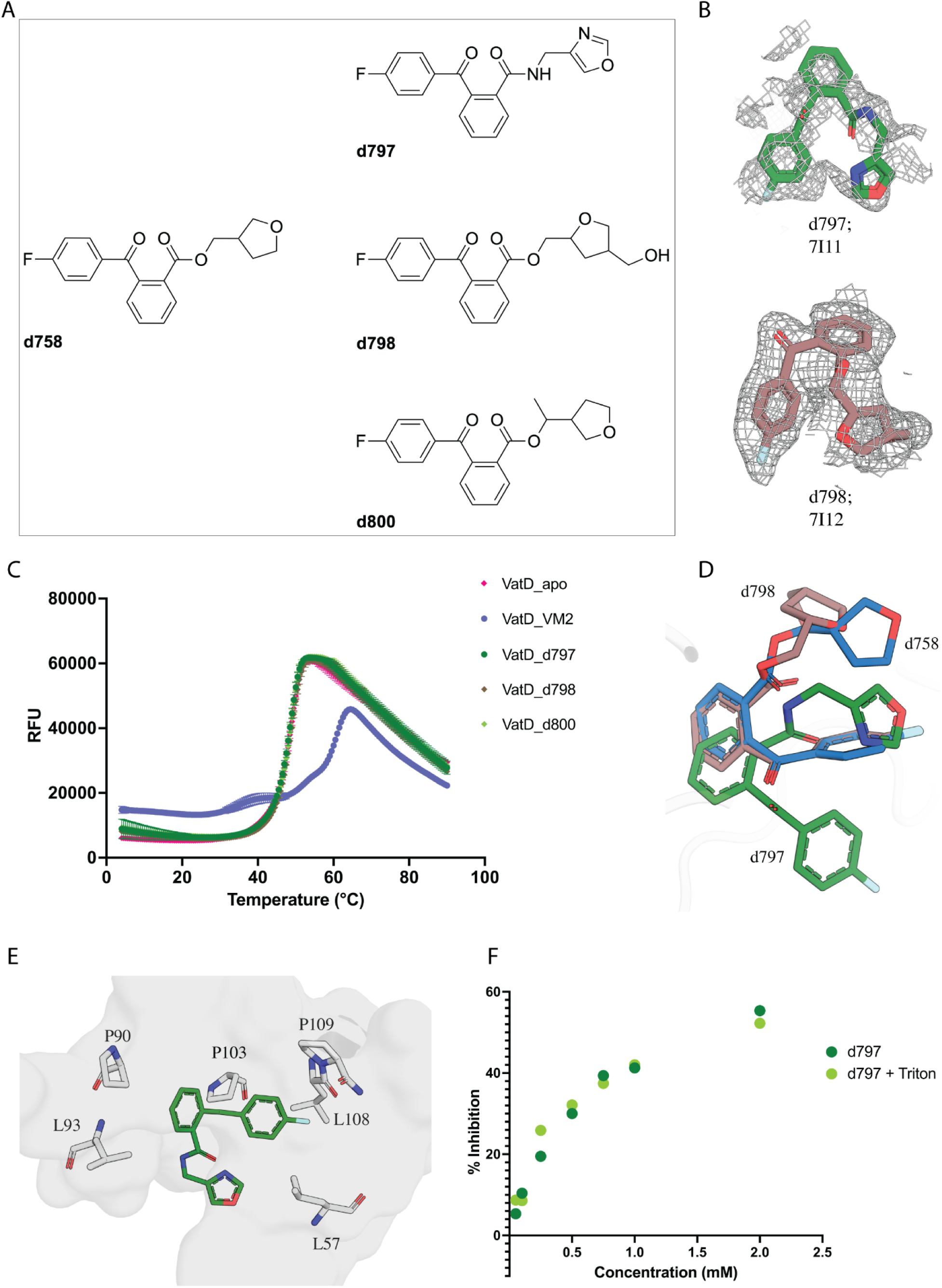
A) Chemical structure of d758 (PDB: 7I0Y) analogs. B) d797 (green, PDB: 7I11) and d798 (brown, PDB: 7I12) PanDDA event density map. C) DSF curve of d797 (green), d798 (brown), and d800 (light green) with VatD-apo (pink) and positive control VatD-VM-2 (purple, PDB: 1KHR). D) Overlap of d797 (green), d798 (brown) on the parent compound d758 (blue). E) d797 hydrophobic interaction with VatD residues. The binding region is shown in surface representation, with the key residues shown in sticks. d797 revealed two new hydrophobic interactions with P103, and P109 residues of VatD. G) The % inhibition graph of d797 in presence (light green) and absence (dark green) of 0.01% Triton-X in the assay. The concentration of d797 ranges from 0.05 mM to 2 mM, in the assay. The x-axis represents the mM concentration of compounds and the y-axis represents the % inhibition.

## Discussion

Streptogramins are a promising class of antibiotics for treating the gram positive bacterial infections. However, the presence of resistance mechanisms, especially Vat enzymes, has significantly reduced their use. Our previous work focused on modifying the streptogramin macrocycle to overcome this resistance mechanism ^17^. To augment this strategy, inhibitors of Vat enzymes could be developed that play an analogous role to β-lactamase inhibitors in combating β-lactam resistance. Our results present several starting points towards this long-term goal.

Our high-throughput crystal soaking experiments with VatD yielded several unexpected insights, providing valuable information for future inhibitor design strategies. The fragment screening result revealed that both the antibiotic binding site and the Ac-CoA sites could be potential targets for inhibitor development. Interestingly, more initial fragment hits were found at the Ac-CoA site. These hits exhibited two distinct receptor binding conformations in the acetyl subsite. However, despite these structural insights, compound d698, which was designed based on hits from the acetyl subsite, failed to exhibit any inhibitory activity and produced an undetectable thermal shift in DSF.

Despite using a fragment library designed to cover a broad chemical space, it is noteworthy that none of the identified fragment hits mimicked the adenosine moiety, with the exception of compounds d545. This suggests a potentially high degree of specificity in the binding pocket for this particular structural feature. While the Ac-CoA site initially appeared promising due to the number of fragment hits, our findings highlight potential obstacles in pursuing this avenue for inhibitor development. Also, understanding that the Ac-CoA is at high concentration in bacteria (20–600 μM) ^25^, competitive inhibition at this site may be challenging to progress for cellular activity.

Our results also indicate that the antibiotic binding site is likely the most promising avenue for inhibitor development. Interestingly, we observed that the compounds derived from the antibiotic-binding fragment demonstrated the ability to bind in the Ac-CoA site. This cross-site binding of compounds highlights the complex nature of ligandability for VatD. By leveraging the overlapping binding sites of d516, d521 and VM-1, we were able to incorporate chemical features of VM-1 into further analogs. This structure-guided approach led to the discovery of compound d758, which demonstrated a significant thermal shift in DSF assay (∼6 °C shift) and showed inhibition at higher concentrations. Although this inhibition was sensitive to detergent, further optimization of d758 resulted in the development of d797, a weak inhibitor that is detergent-independent. Interestingly, d797 adopts a different binding pose than the parent compounds, but still has spatial overlap with the antibiotic. As is often seen in fragment-based drug design, these early compounds do not unambiguously pass all the checks of our assays (crystallography, DSF shift, and catalytic assay). However, these results collectively validate the ligandability of this site for the development of future competitive inhibitors with improved affinity and specificity for VatD. Future development of more potent compounds would enable combination therapies that can potentially resurrect the clinical use of streptogramins.

## Methods

### Cloning, protein expression, and purification

Full length VatD was cloned into a pET28a vector with a N-terminal 6x His tag followed by a tobacco etch virus (TEV) protease cleavage site. For expression, the VatD-pET28a plasmid was transformed into BL21-RIPL competent cells. The primary culture was grown overnight in LB (lysogeny broth) broth at 37 °C supplemented with Ampicillin (conc) and Chloroamphenicol (conc). A secondary culture was grown in TB (Terrific broth, 1 Liter) broth at 37 °C until an optical density of 0.8 was reached. Before protein expression, the culture was incubated at 4 °C to cool down for 20 mins. The culture was then induced with 1 mM IPTG (isopropyl b-d-1-thiogalactopyranoside) and were allowed to grow at 18 °C, shaking for 14 hours. The cells were harvested by centrifugation at 4000 g and stored at −80 °C until use.

The cell pellet was resuspended in the buffer A (50 mM HEPES, 500 mM NaCl, 10 mM imidazole, 1 mM TCEP (tris(2-carboxyethyl)phosphine) supplemented with a protease inhibitor tablet and TurboNuclease (5 U/ml; Sigma-Aldrich, T4330)]. The cells were lysed by sonication and the lysate was clarified by centrifugation at 18,000 rpm for 40 mins. The supernatant was then filtered using a 0.2 µm filter (Millipore) and applied to a 5 ml His-Trap HP column(Cytiva, 17524802). The column was then washed with buffer A, followed by buffer B (Buffer A + 50 mM Imidazole). The protein was eluted in buffer A containing 500 mM Imidazole. The eluted protein was incubated with TEV protease (1:10 protein:enzyme) exchanged into a TEV reaction buffer at 4 °C for 16 hours. The cleaved protein was centrifuged at 4000 rpm, 4 °C for 15 mins and the supernatant was loaded onto a pre-equilibriated 5 ml His-Trap HP column to separate the cleaved protein from TEV protease and any uncleaved protein. The cleaved protein was eluted using buffer A and concentrated to 2.5 ml using a 10 kDa molecular weight cut off Amicon concentrator and injected onto a pre-equilibriated Superdex 200 120 ml column (buffer: 50 mM HEPES, 150 mM NaCl, 1 mM TCEP (tris(2-carboxyethyl)phosphine). Eluted fractions were concentrated to 20 mg/ml and flash frozen in liquid nitrogen until further use.

### Crystallization and fragment soaking

VatD crystals were grown using sitting drop vapor diffusion method at +20 °C in a reservoir solution containing 0.1M Tris, 25% Peg 1000, pH 8.0. The crystallization drops were setup in a 3-well MRC plate (SWISSCI, 3W96T-UVP) with 200 nl of purified protein and 200 nl of reservoir solution. Crystals appeared withing 2 days and take ∼4-5 days to grow to the maximum size before soaking with fragments.

Before soaking the fragments, the crystals were tested for DMSO tolerance with 5%, 10% and 20% DMSO which showed that crystals dissolved/cracked with 20% DMSO. However, crystals soaked with 5% and 10% DMSO remained intact without any change in morphology or resolution. 37 crystals were soaked with 10% DMSO only, classified as the apo crystals. Two fragment libraries were used to soak the crystals: the Enamine Essential fragment library containing 320 fragments and the UCSF_91 library contating 91 fragments. The crystals were soaked with 10% DMSO or 10% fragment using an acoustic dispenser with an Echo 650 liquid handler (Labcyte) and the plates were incubated at +20 °C for 2-4 hours. The crystals were harvested after the incubation without using any cryoprotectant. The hits from the first round of soaking were used to generate lead like compounds. These compounds were ordered from the Enamine store (REAL DATABASE).

### Crystal data collection, processing, and fragment modelling

The diffraction data of VatD soaked with DMSO and fragments were collected at 100 K on the ALS synchrotron beamline 8.3.1 and SSRL beamline 12-1, and 12-2. The data collection parameters are mentioned in supplementary excel file. The datasets were indexed, integrated and scaled using XDS ^26^ and were merged using AIMLESS ^26,27^. The apo structure (soaked in 10% DMSO) was solved using the expert mode-molecular replacement in CCP4 with reference structure of VatD (PDB: 1MR7) ^28^. The refinement was carried out in Phenix using phenix.refine^29^ with iterative rounds of manual model building in Coot ^30^.

The fragment soaked datasets were processed using the DIMPLE pipeline using the apo structure (soaked in 10% DMSO) as the reference molecule ^31^. PanDDA analysis was performed to identify the fragment density ^20^. A total of 33 DMSO only datasets were collected which served as the background electron density maps for PanDDA. The ligand restraints were generated using *phenix.elbow* ^32^.

### Differential Scanning Fluorimetry (DSF)

Differential Scanning Fluorimetry (DSF) was performed to assess the thermal stability of the protein in the presence of Ac-CoA, VM-2 and elaboration compounds. The protein was used at a final concentration of 5 µM. The compounds were prepared at 10 mM in the protein buffer and diluted to 2 mM in the reaction mixture. The final DMSO concentration in the assay was kept at 2%. SYPRO Orange dye (Sigma-Aldrich) was used as the fluorescent reporter at a 4X final concentration. The samples were prepared in 96-well semi-skirt 0.1ml PCR plate (1402-8990, USA scientific) with triplicate wells for each condition. The total reaction volume for each condition was 25 µl. The plate was sealed with optical adhesive film, and spinned down for 1 min. A thermal gradient was applied from 25°C to 95°C at a rate of 1°C per minute, and fluorescence was recorded at each temperature increment. The fluorescence intensity was monitored as a function of temperature. The melting temperature (Tm) was calculated using the web application DSF world (https://bio.tools/dsfworld) and the graphs were plotted in the prism software.

### Kinetic assay

#### VatD Acetylation Kinetics with Respect to VM-2 and AcCoA

The reaction rate of VatD was determined using a colorimetric assays measuring the formation of coenzyme A (CoA), a by-product of VatD-catalyzed acetylation of streptogramins with acetyl-CoA (Ac-CoA). The reaction of 5,5’-dithiobis-(2-nitrobenzoic acid) (DTNB) with the free sulfhydryl of CoA produces 2-nitro-5-thiobenzoate (TNB), the absorbance of which was measured at 415 nm in a Spark Multimode Microplate Reader with 384-well clear polystyrene flat-bottom NBS plates (Corning 3544). The kinetic parameters of VatD acetylation were determined with respect to varied concentrations of Virginiamycin 2 (VM-2) and Ac-CoA.

To measure reaction rate with respect to VM-2, dilutions of VM-2 were prepared in DMSO for a final concentration of 0.0025-0.25 mM VM-2 and 5% DMSO in each well. The assay composition consisted of 0.5 mM DTNB and 1 mM AcCoA in 100 mM NaCl and 100 mM HEPES pH 7.8 buffer.

To measure reaction rate with respect to AcCoA, dilutions of AcCoA were prepared in nuclease free water for final concentrations ranging from 0.005-0.5 mM AcCoA in each well. The assay composition was identical to that previously described, except 1 mM AcCoA was exchanged for 1 mM VM-2 in each well. For all kinetic reactions, VatD was spiked into the reaction mixture at a final concentration of 29 nM in each well, and readings were performed at room temperature in duplicate.

A standard curve was prepared to quantify CoA production. The standard curve assay was composed of 0.5 mM DTNB in 100 mM NaCl and 100 mM HEPES pH 7.8 buffer with a dilution series of CoA at a final concentration of 0 to 0.8 mM in each well. Initial reaction rates were determined using Excel by fitting the data to a linear regression model within the linear range of each reaction curve. The standard curve was used to convert these rates from Absorbance/second to CoA (mM)/second/enzyme (nM). Kinetic parameters were derived by fitting the initial reaction rates at varied concentrations of either VM-2 or Ac-CoA to the Michaelis-Menten model using GraphPad Prism version 10.0.0 for Mac (GraphPad Software, Boston, Massachusetts USA, www.graphpad.com).

#### Inhibition Dose-Response with Fragments

To measure the dose-response of each potential inhibitor, fragments were prepared from 100 mM stock solutions in DMSO and diluted to concentrations of 2 mM, 1 mM, 0.75 mM in each well. Selected fragments which showed inhibitions at higher concentrations were further diluted to 0.5 mM, 0.25 mM, 0.1 mM, and 0.05 mM in each well to expand the measured range of dose-dependent inhibition. The final concentration of DMSO in each well was 5%.

In addition to the inclusion of fragments, the assay composition of these dose-response curves was altered to contain either VM-2 or Ac-CoA at their respective Km concentrations. For fragments binding at the antibiotic-binding site, their dose-response assay contained a fixed concentration of 0.05 mM VM-2 and was otherwise identical to that measuring kinetic parameters with respect to VM-2. Similarly, aside from the addition of fragments, the assay composition for testing fragments binding at the Ac-CoA-binding site contained a fixed concentration of 0.15 mM AcCoA and was otherwise identical to that measuring kinetic parameters with respect to AcCoA. Relative inhibition was measured by comparing the initial reaction rate at either 0.05 mM VM-2 or 0.15 mM Ac-CoA in the presence and absence of fragments.

#### VatD Acetylation Kinetics with Detergent

To distinguish aggregation-based inhibition from that of competitive fragment binding, dose-response assays of selected fragments were repeated with the addition of 0.01% Triton-X vol/vol. To determine the impact of 0.01% Triton-X vol/vol on the enzyme or other assay components, we repeated the kinetic assays without fragment. The assay buffer was incubated with 0.01% Triton-X vol/vol for 5 minutes prior to the addition of enzyme.

## Supporting information

Supplemental X-ray Data

## Acknowledgements

J.S.F is supported by NIH GM145238. The diffraction data of structures reported in this work was collected at the ALS (8.3.1) and SSRL (9-2 and 12-2) beamlines. Use of the SSRL, SLAC National Accelerator Laboratory, is supported by the U.S. Department of Energy, Office of Science, Office of Basic Energy Sciences under Contract No. DE-AC02-76SF00515. The SSRL Structural Molecular Biology Program is supported by the DOE Office of Biological and Environmental Research, and by the National Institutes of Health, National Institute of General Medical Sciences (P30GM133894). The ALS, a U.S. DOE Office of Science User Facility under contract no. DE-AC02-05CH11231, is supported in part by the ALS-ENABLE program funded by the NIH, National Institute of General Medical Sciences, grant P30GM124169.

## Author contributions

PA: Conceptualization, Methodology, Investigation, Formal analysis, Writing - original draft, and review & editing. JSF: Conceptualization, Funding acquisition, Writing - original draft, and review & editing. SL: Investigation (enzyme kinetics assay). IS: Conceptualization, review & editing, CM: review & editing.

## Data Availability

The fragment-bound datasets have been deposited in RCSB-PDB under group deposition: G_1002332. The individual PDB ID for each fragment-bound dataset is given in supplementary table 1. The background DMSO datasets used in this study are uploaded on Zenodo (DOI: 10.5281/zenodo.14775497)

## Competing interests

J.S.F. is a consultant to, shareholder of, and receives sponsored research support from Relay Therapeutics.

## Supplementary Information

**Supplementary Figure 1:**
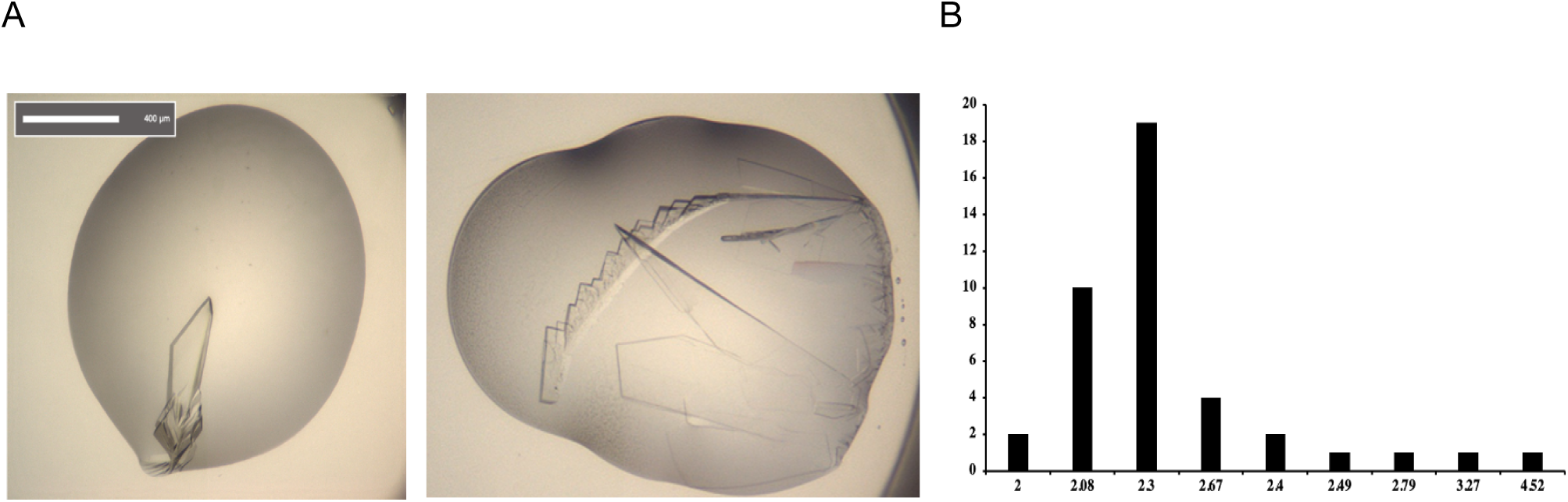
A) Vat D crystals grown in 0.1M Tris pH 8.5, 25% w/v Peg 1000 at 20 °C. B) Vat D crystals grown in 0.2M NH_4_Cl, 17% Peg 3350 B) Histogram showing the resolution range of VatD crystals soaked with 5 % DMSO (black) for 4 hours.

**Supplementary Figure 2:**
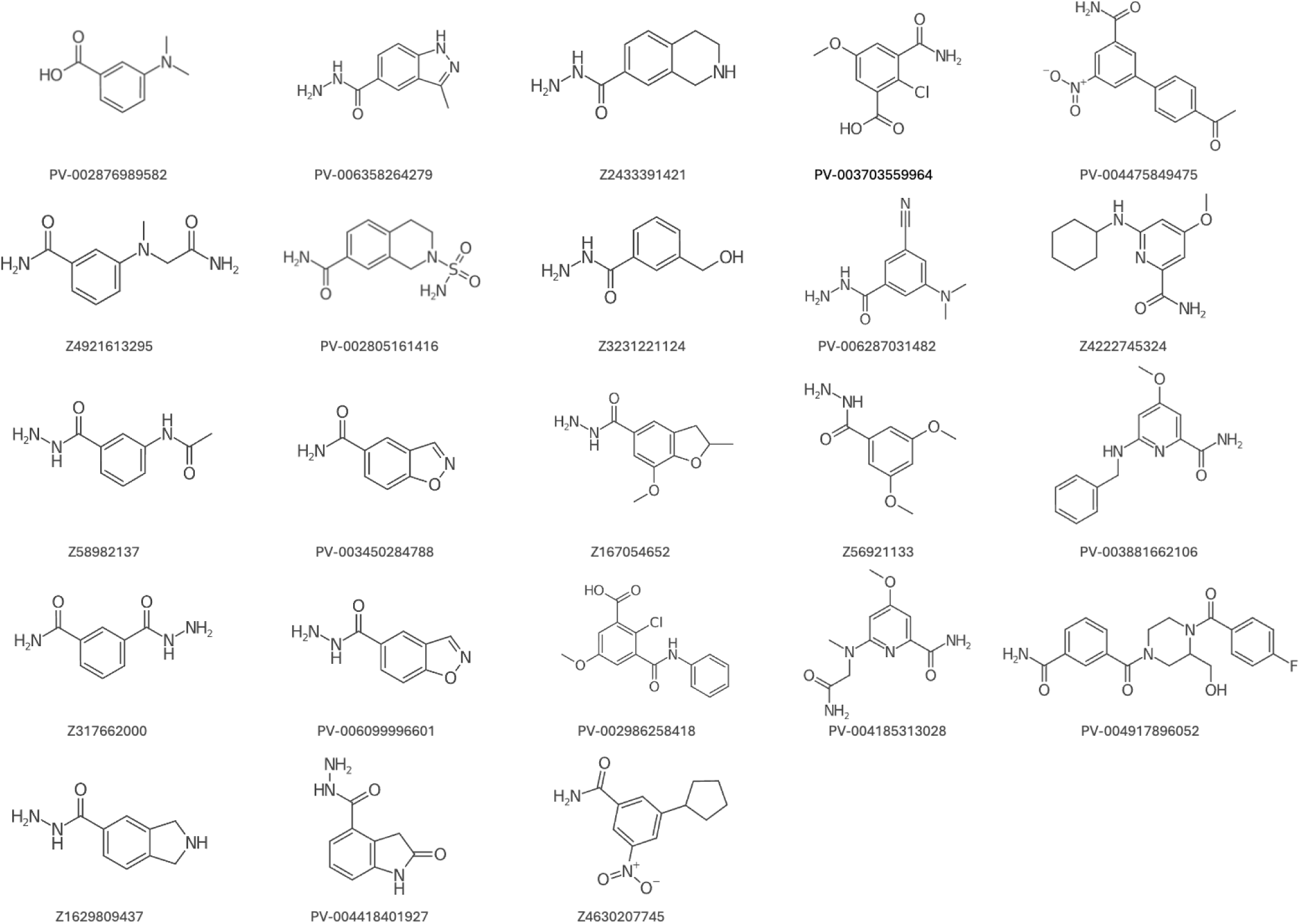
Elaboration compounds of d384.

**Supplementary Figure 3:**
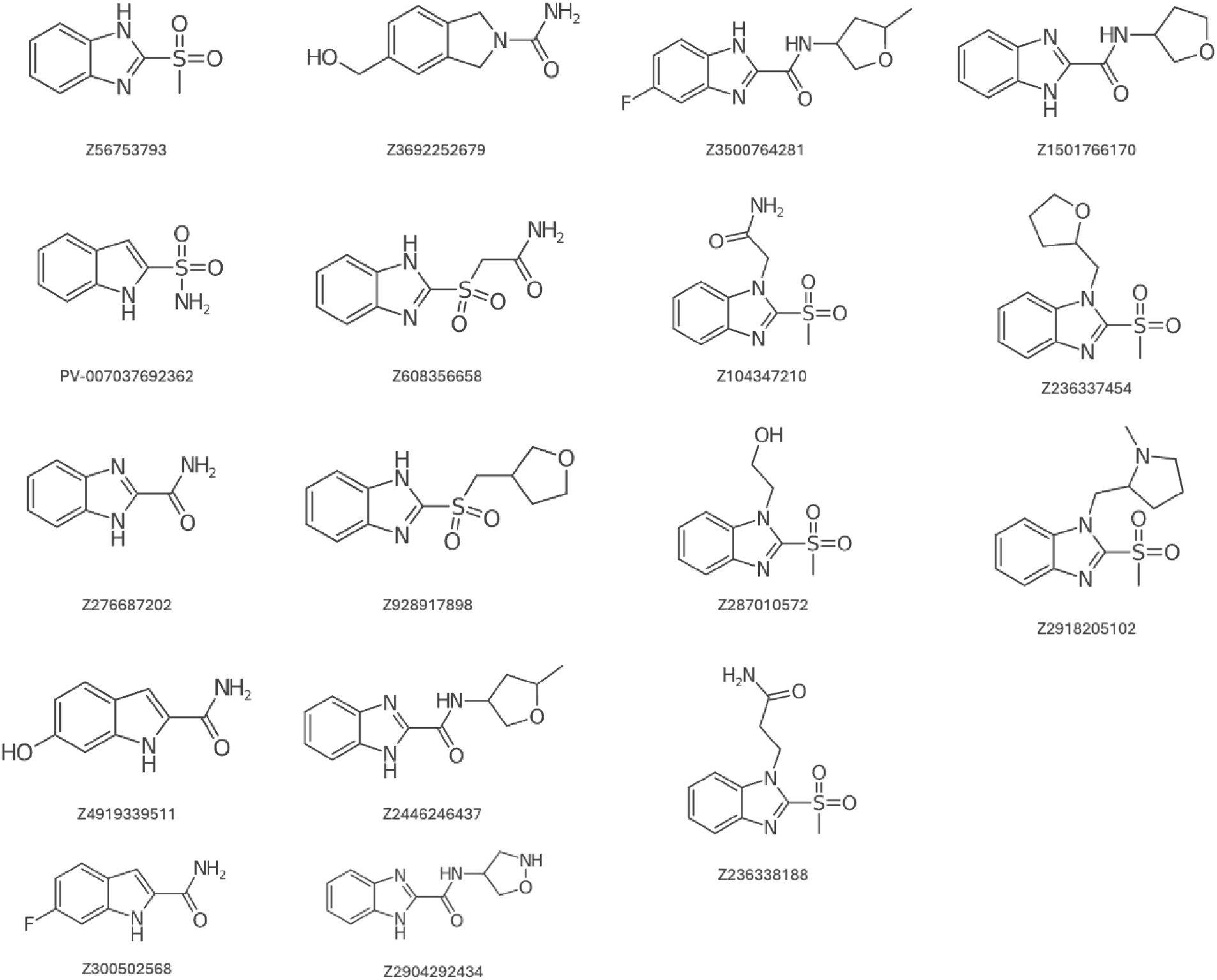
Elaboration compounds of d521

**Supplementary Figure 4:**
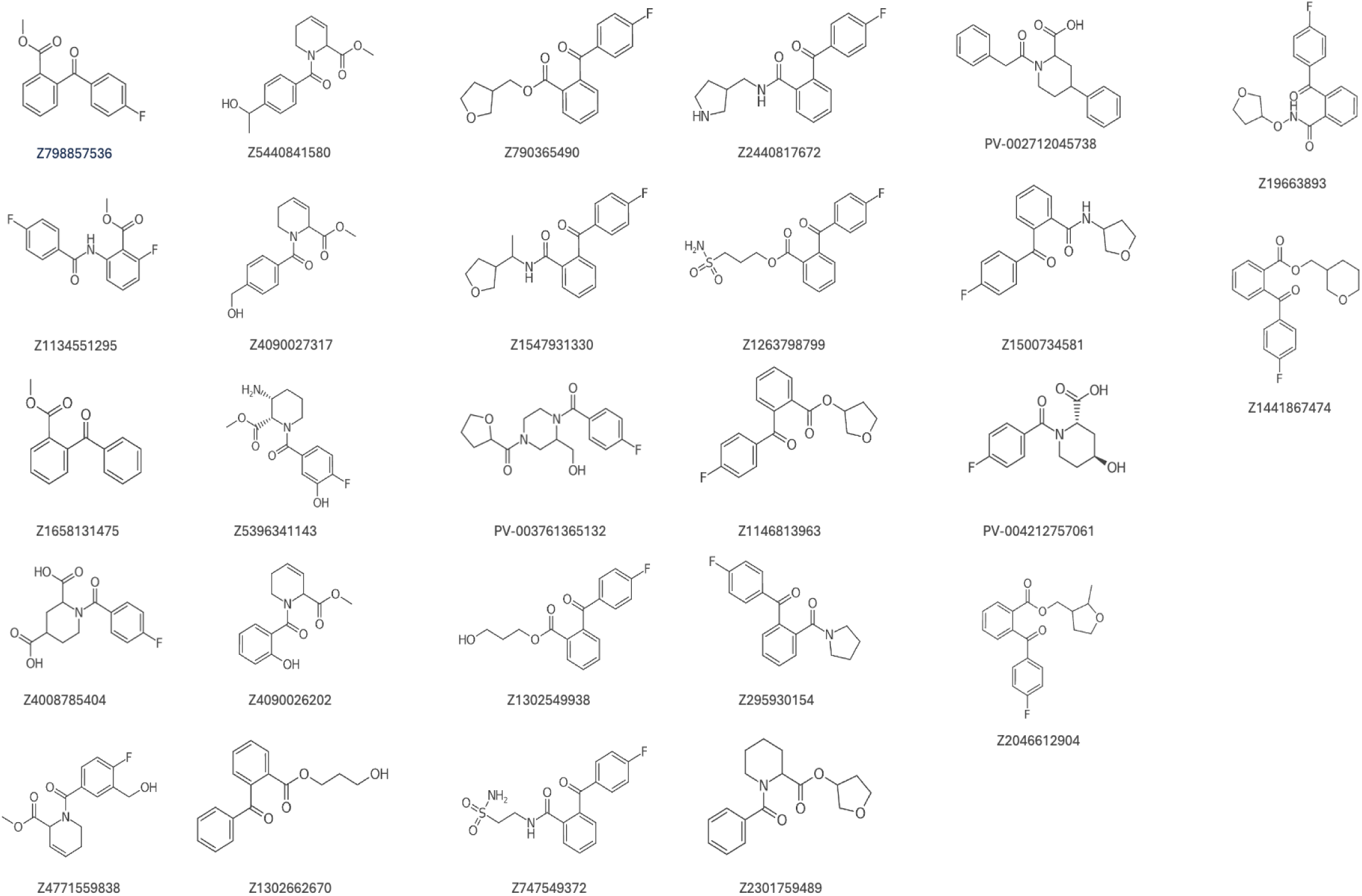
Elaboration compounds of d516

**Supplementary Figure 5:**
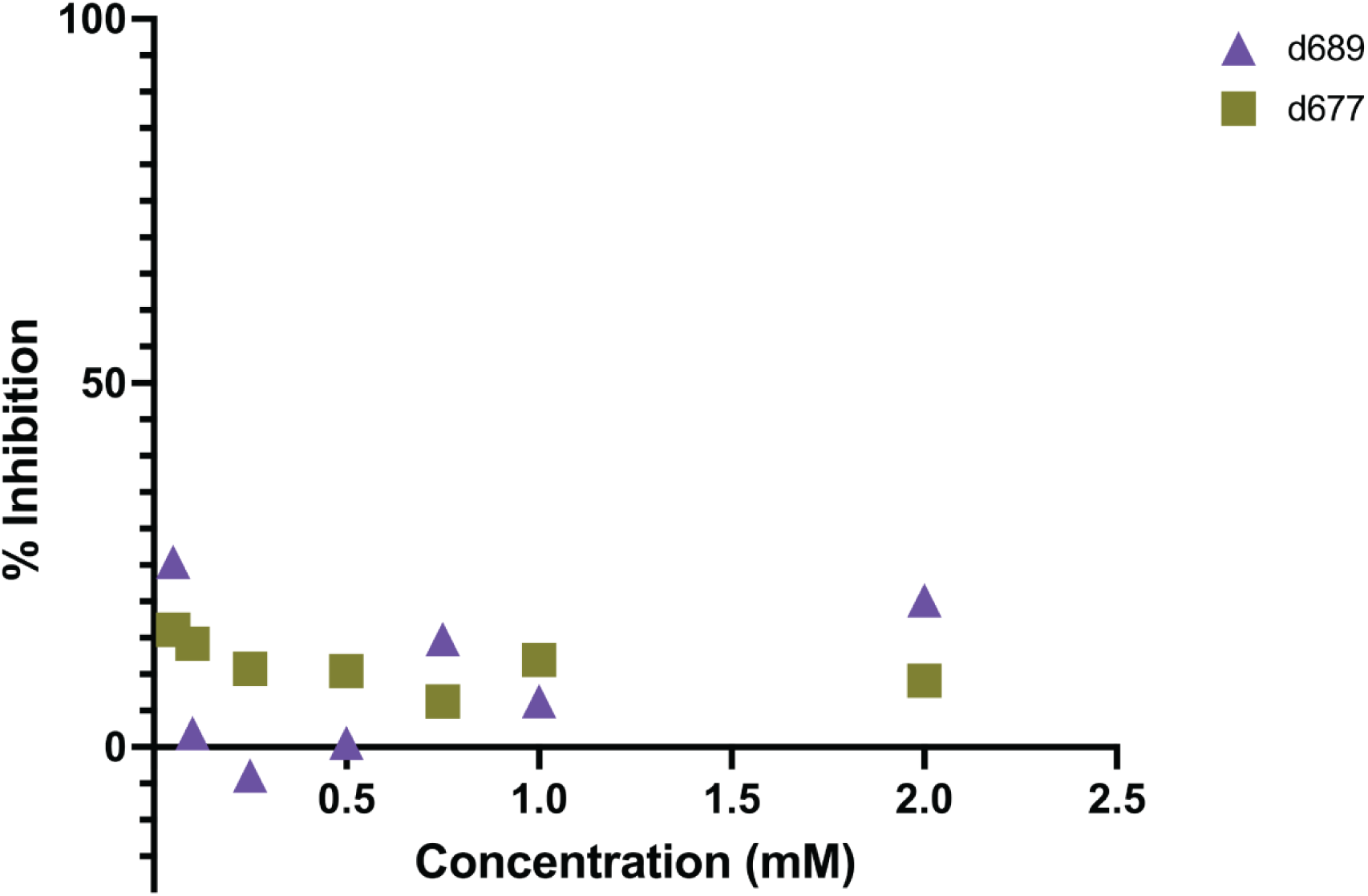
Inhibition curve of d677 and d689

**Supplementary Figure 6:**
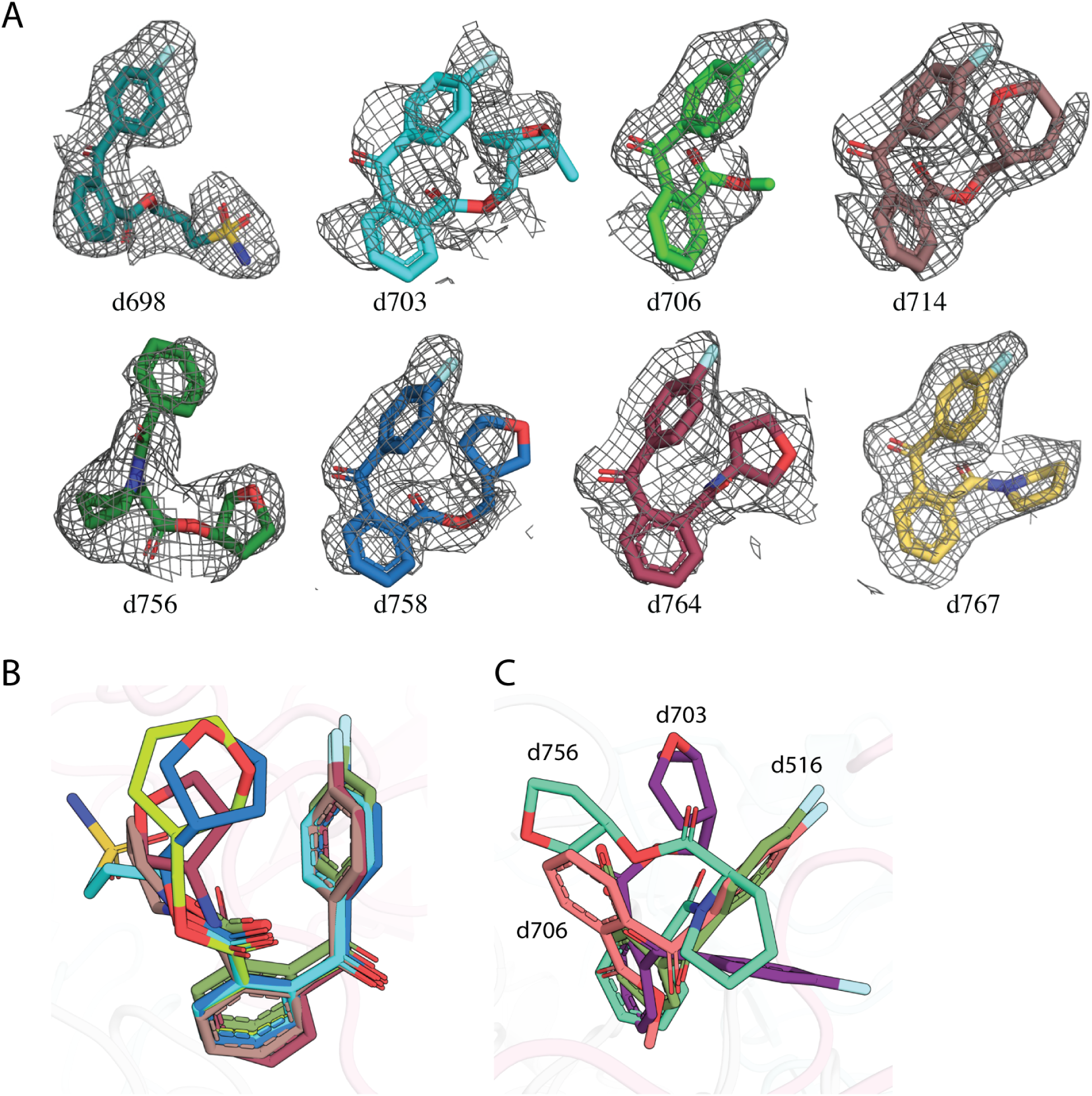
A) PanDDA event density maps of d516 (PDB: 7I0F) analogs that bind in the antibiotic site. B) Compounds showing complete overlap with the parent fragment d516. C) Compounds d703 (PDB: 7I0S), d706 (PDB: 7I0U), and d756 (PDB: 7I0X) not overlapping with the parent fragment d516.

**Supplementary Figure 7:**
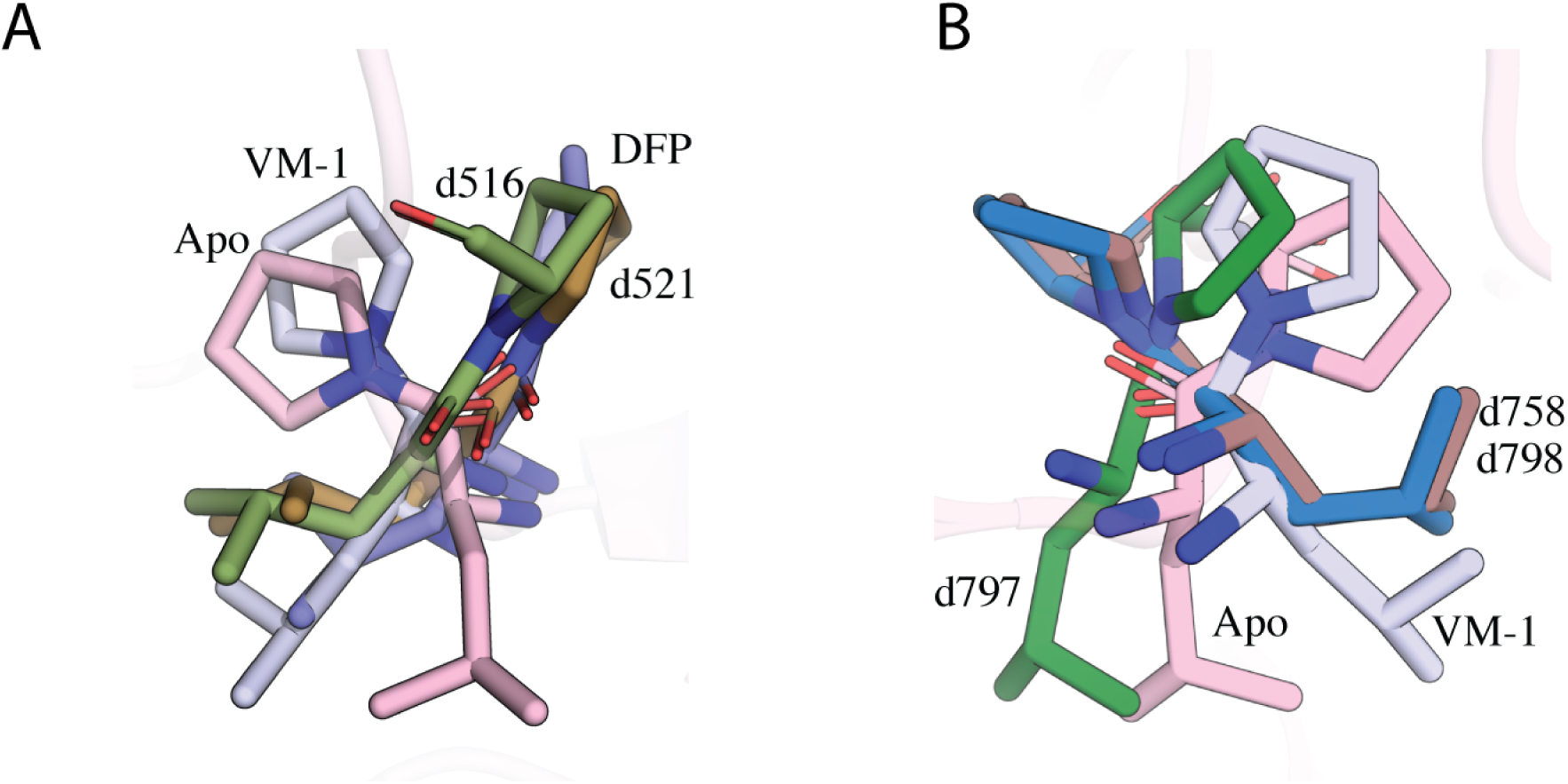
A) Conformational flip in L108 and P109 residues in d516 (PDB: 7I0F) and d521 (PDB: 7I0H). B) Conformational flip in L108 and P109 residues of d516 analogs d758 (PDB: 7I0Y), d797 (PDB: 7I11), and d798 (PDB: 7I12).

**Supplementary Figure 8:**
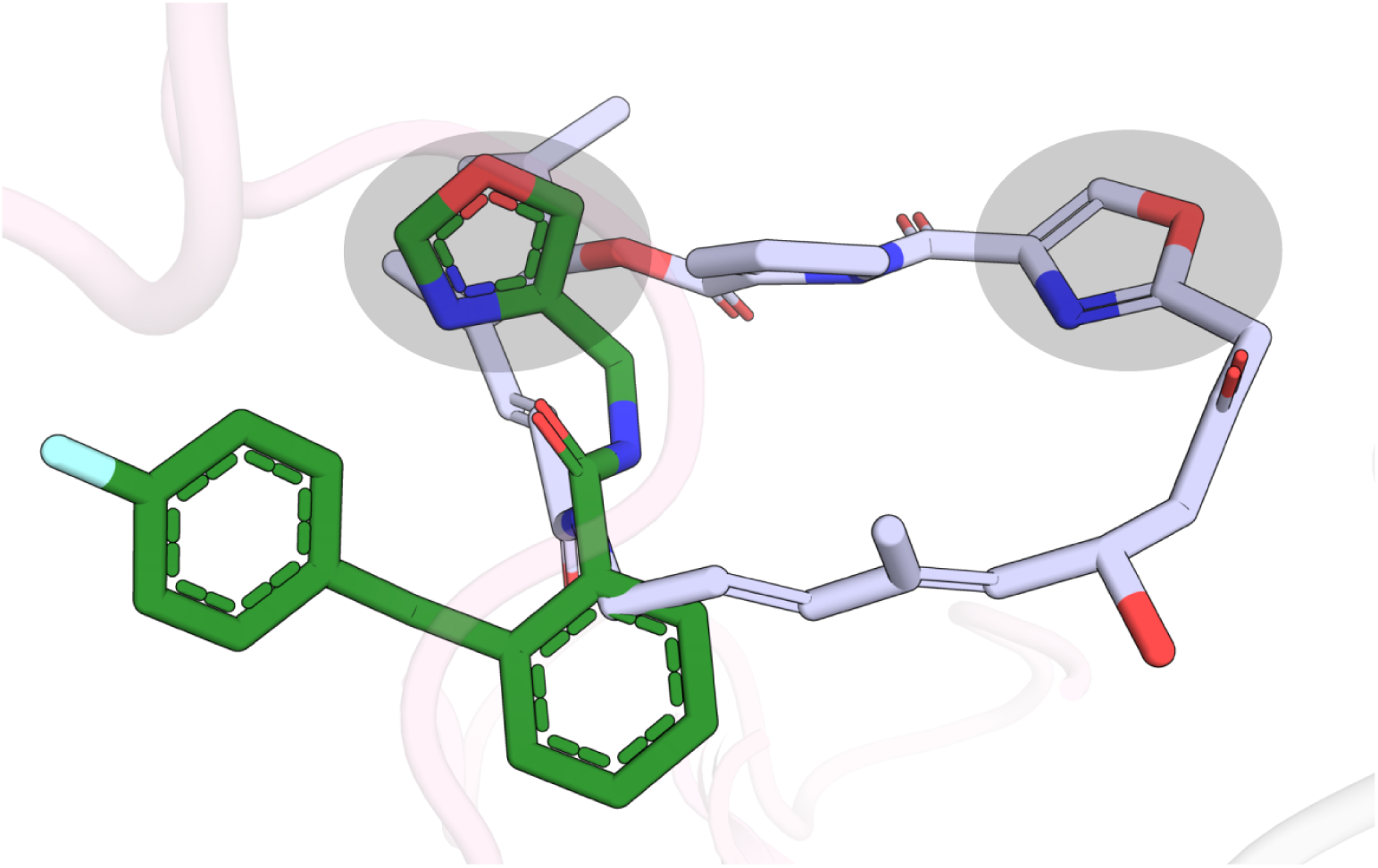
Placement of furan ring in d797 (PDB: 7I11) and VM-1 (PDB: 1KHR) bound VatD complexes. Grey shaded region highlights the furan ring in both compounds.

**SI Table 1:**

| Model | Pdb ID |
| --- | --- |
| d116 | 7HZV |
| d167 | 7HZW |
| d193 | 7HZX |
| d229 | 7HZY |
| d232 | 7HZZ |
| d275 | 7I00 |
| d287 | 7I01 |
| d330 | 7I02 |
| d374 | 7I03 |
| d384 | 7I04 |
| d397 | 7I05 |
| d398 | 7I06 |
| d425 | 7I07 |
| d429 | 7I08 |
| d436 | 7I09 |
| d438 | 7I0A |
| d440 | 7I0B |
| d474 | 7I0C |
| d499 | 7I0D |
| d501 | 7I0E |
| d516 | 7I0F |
| d519 | 7I0G |
| d521 | 7I0H |
| d537 | 7I0I |
| d538 | 7I0J |
| d545 | 7I0K |
| d560 | 7I0L |
| d619 | 7I0M |
| d626 | 7I0N |
| d666 | 7I0O |
| d677 | 7I0P |
| d689 | 7I0Q |
| d698 | 7I0R |
| d703 | 7I0S |
| d705 | 7I0T |
| d706 | 7I0U |
| d714 | 7I0V |
| d724 | 7I0W |
| d756 | 7I0X |
| d758 | 7I0Y |
| d764 | 7I0Z |
| d767 | 7I10 |
| d797 | 7I11 |
| d798 | 7I12 |

## Notes

https://10.5281/zenodo.14775497

### Summary of Updates

Slight reformatting, fixing names, adding funding.

